# Taurine/chenodeoxycholic acid ratio as a circulating biomarker of insidious vitamin B_12_ deficiency in humans

**DOI:** 10.1101/2024.02.15.580499

**Authors:** Madhu Baghel, Sting L. Shi, Himani Patel, Vidya Velagapudi, Abdullah Mahmood Ali, Vijay K. Yadav

## Abstract

Deficiency of vitamin B_12_ (B_12_), an essential water-soluble vitamin, leads to irreversible neurological damage, osteoporosis, cardiovascular diseases, and anemia. Clinical tests to detect B_12_deficiency lack specificity and sensitivity. B_12_ deficiency is thus insidious because progressive decline in organ functions may go unnoticed until the damage is advanced or irreversible. Here, using targeted unbiased metabolomic profiling in the sera of B_12_-deficient versus control individuals, we set out to identify biomarker(s) of B_12_ deficiency. Metabolomic profiling identified 77 metabolites, and Partial least squares discriminant-analysis (PLS-DA) and hierarchical clustering analysis (HCA) showed a differential abundance in B_12_-deficient sera of taurine, xanthine, hypoxanthine, chenodeoxycholic acid, neopterin, and glycocholic acid. Random forest (RF) multivariate analysis identified a taurine/chenodeoxycholic acid ratio, with an AUC score of 1, to be the best biomarker to predict B_12_ deficiency. Mechanistically, B_12_ deficiency reshaped the transcriptomic and metabolomic landscape of the cell identifying a downregulation of methionine, taurine, urea cycle, and nucleotide metabolism, and an upregulation of Krebs cycle. Thus, we propose taurine/chenodeoxycholic acid ratio in serum as a potential biomarker of B_12_ deficiency in humans and elucidate cellular metabolic pathways regulated by B_12_ deficiency.

## INTRODUCTION

Vitamin B_12_ (B_12_) is an essential water-soluble vitamin derived from animal-based diets that regulates a multitude of cellular processes in humans such as one-carbon metabolism and Krebs cycle.(1–4) The absorption of dietary B_12_ requires gastric intrinsic factor (GIF), a stomach-specific protein.(4) Gif binds to B_12_ in the small intestine forming the GIF-B_12_ complex. This complex is endocytosed by the intestinal epithelial cells and B_12_ is released into the bloodstream.(4) In the bloodstream, B_12_ binds to the protein transcobalamin 2, which then carries it to the liver, the primary storage and recycling organ for B_12_ in mammals.(5) Once acquired, humans, for instance, can recycle B_12_ to maintain B_12_-dependent cellular processes for up to a decade.(2) In the cells, B_12_–derivatives function as cofactors for only two known enzymes: methylmalonyl-CoA mutase and methionine synthase, and through them affect a variety of downstream metabolic pathways such as Krebs cycle, amino acid synthesis, and DNA and histone methylation. (1, 6) In humans, decreased production of functional GIF protein or non-consumption of animal products causes B_12_ deficiency and results in various abnormalities, such as anemia, osteoporosis, and cognitive defects. (7–10)

In clinical practice, the diagnosis of B_12_ deficiency is typically established by the measurement of serum cobalamin (Cbl) levels.(11) Although B_12_ deficiency can be reflected by elevated methylmalonic acid (MMA) and homocysteine (Hcy) levels, these tests are not routinely used unless the initial Cbl levels are equivocal because MMA and Hcy can be elevated in conditions independent of B_12_ levels.(12–16) Despite the importance of B_12_ and its association with many physiological functions, many issues remain unresolved in the diagnosis of B_12_ deficiency, leading to poor diagnosis and irreversible consequences on the body.(17, 18) First, B_12_ is a very stable molecule and because 95-97% of B_12_ is stored in the liver, its serum levels do not accurately reflect its actual functional levels i.e., the amount of B_12_ required for maintaining body functions.(19) Second, the cost of measurement of B_12_ in patient samples, despite being not able to accurately predict a B_12_-deficient state, remains high and therefore is not the first line of measurement; clinicians measure B_12_ only when a patient presents signs of B_12_ deficiency such as anemia to confirm a deficient state. (20–23) These facts necessitate the need to identify molecules regulated by B_12_, which can provide a functional readout of B_12_ deficiency in humans.

We recently created a transgenic mouse model of B_12_ deficiency by deleting the gene essential for B_12_ absorption from the gut, *Gif*, to understand the molecular consequences of B_12_ deficiency. These studies led to the identification that B_12_ stored in the liver regulates the production of taurine. Taurine is a semi-essential micronutrient that has recently been shown to be a driver of aging as its supplementation increases healthy lifespan in diverse species from worms to mice, and low taurine levels are associated with poor health in aged humans(24). In the B_12_ mode of action, taurine plays an important role as the reversal of taurine deficiency through daily oral taurine administration was shown to fully rescue the consequences of B_12_ deficiency(25). More importantly, the targeted metabolomics analysis of liver tissue collected from control and B_12_-deficient mice showed changes in a multitude of metabolites besides taurine that are secreted from cells and could be detected in the serum(25). These studies suggested a plausible and testable hypothesis that certain metabolites or sets of metabolites may exist which could serve as a readout of, difficult to detect, B_12_-deficient state in humans.

The present study was initiated to test the above hypothesis by performing a metabolomic analysis on serum samples collected from control and B_12_-deficient individuals to identify which factor(s) could serve as a biomarker of B_12_-deficient state. Results showed that serum levels of certain metabolites such as taurine, xanthine and hypoxanthine were dramatically downregulated in the B_12_-deficient individuals. Using various downstream analyses, we suggest that taurine in conjugation with chenodeoxycholic acid can serve as a biomarker of B_12_-deficient state in humans. Furthermore, using mouse B_12_-deficient tissues, we elucidate how despite only needed for 2 enzyme functions, B_12_ deficiency alters the metabolic and transcriptomic landscape in the cells, which will facilitate advances in further understanding biology of B_12_.

## RESULTS

### Study population, sample classification, acquisition, pre-processing, and normalization of metabolomic data

A schematic diagram illustrating different steps of this study is presented in **Figure 1**. The samples utilized in this study are from the Kuopio Ischaemic Heart Disease Risk Factor (KIHD) study aimed at identifying the risk factors for coronary heart diseases, atherosclerosis, and other related conditions in the Eastern Finnish population.(26) Sera were classified in accordance with internationally established criterion into control subjects (n=13) with B_12_ levels >250 pmol/L, and into deficient subjects (n=8) with B_12_ levels <150 pmol/L.(1, 11, 17, 27) Samples were randomized before metabolite extraction and quantified using a ACQUITY UPLC-MS/MS system. Ninety-four metabolites could be detected in the sera, out of which 77 that passed quality control were selected for further downstream analysis. Imputation of one missing value with the minimum value in that cohort was done, and data was pre-processed by generalized log transformation (glog) and auto-scaling of metabolite concentration peaks in each sample to represent uniform distribution.

**FIGURE 1:**
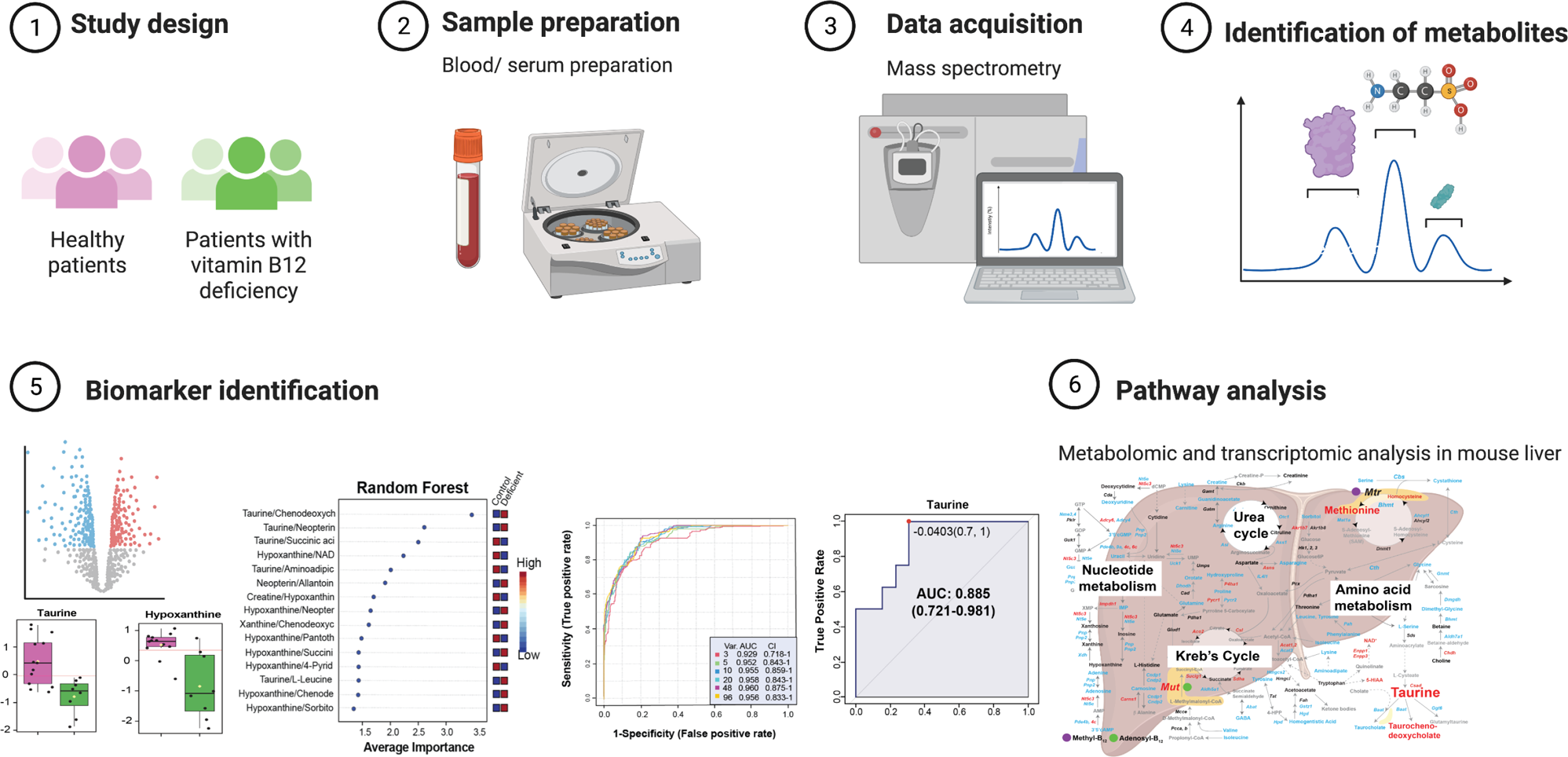
Study population, sample classification, acquisition, pre-processing, and normalization of metabolomic data. Schematic diagram illustrating the steps for metabolomic analysis of serum samples from B_12_-deficient (B_12_ levels <150 pmol/L) versus the healthy control group. (1) In this study, 8 and 13 subjects were grouped in B_12_-deficient and control groups (age- and gender-matched), respectively, (2) blood samples were collected and processed, (3) metabolomics data was acquired from serum samples using ACQUITY UPLC-MS/MS system (Waters Corporation, Milford, MA, USA), data was pre-processed and analyzed using MetaboAnalyst 5.0 to identify (4) differentially expressed metabolites between 2 study groups, (5) serum metabolic biomarker for Vitamin B_12_ deficiency followed by (6) pathway analysis.

### Identification of differentially expressed serum metabolites following B_12_ deficiency

We first performed a principal component analysis (PCA), an unsupervised multivariate analysis, to group/classify samples without any consideration of prior classification to detect any outliers in the two cohorts. The principal component 1 (PC1) accounted for 22.6% of the variance and PC2 accounted for 13.6% of the variance (**Figure 2A**). To identify differential concentration of each metabolite between the control and B_12_-deficient groups, we calculated the mean fold change and performed t-tests to compare the mean of each metabolite. A metabolite was considered significantly different between each group when the value of p ≤ 0.05 and log2 fold change ±0.5. In the colvano plot the 3 blue dots in the upper left and 3 red dots in upper right quadrants represent the most significantly altered metabolites in B_12_-deficient subjects compared to that in controls (**Figure 2B**). A hierarchical clustering analysis (HCA) of the metabolomic data using the top 3 downregulated and top 3 upregulated metabolites showed well-defined clustering of thirteen healthy subjects (pink, left cluster) versus eight B_12_-deficient subjects (green, right cluster) (**Figure 2C**). The control group showed high abundance (shades of red colour) of taurine, hypoxanthine and xanthine compared to the B_12_-deficient group, whereas the abundance of glycocholic acid, neopterin and chenodeoxycholic acid was significantly higher in the B_12_-deficient group as compared to healthy controls (**Figure 2C**). Following the identification of differentially expressed metabolites (DEMs), we did Metabolite Set Enrichment Analysis (MSEA) and Metabolomic Pathway Analysis (MetPA) to determine the metabolic pathways that are associated with differences in the abundance of identified metabolites, and perturbations of which is associated with the B_12_ deficiency. The MSEA classified the 77 DEMs into 50 different metabolic pathways (**Figure 2D**) that include divergent cellular metabolism pathways such as bile acid biosynthesis, amino acid biosynthesis, glucose metabolism, and nucleic acid synthesis, which are listed in the order of descending fold enrichment (**Figure 2D**). Out of the 50 listed pathways, the taurine and hypotaurine metabolism pathway was the most enriched pathway with highest fold enrichment value (-logP value ∼6). MetPA results revealed that taurine and hypotaurine metabolism pathway had the highest pathway impact value between the controls and B_12_-deficient subjects, further validating the importance of this pathway (**Figure 2E**).

**FIGURE 2:**
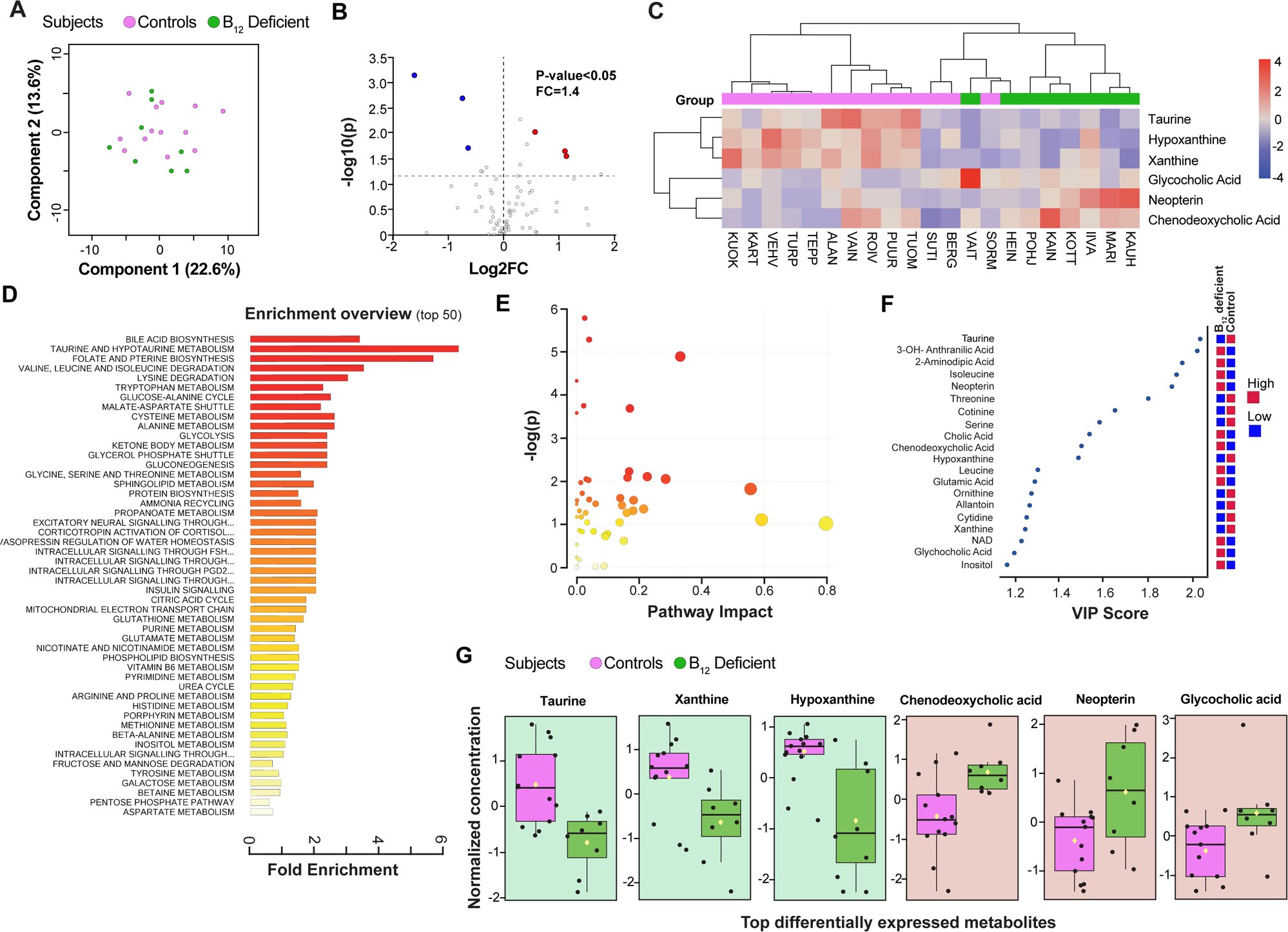
Identification of differentially expressed serum metabolites following B_12_ deficiency. (A) Unsupervised multivariate PCA plot showing the spread of control (pink dots) versus B_12_-deficient (green dots) cohort based on the serum metabolic profile. The horizontal and vertical coordinates are the first and second principal components, respectively. Each dot represents a sample. (B) Volcano plot showing six (blue and red dots) most significant differentially expressed metabolites between the B_12_-deficient patients versus controls, with a p-value < 0.05 and a log2 fold change ±0.5. X-axis corresponds to log2(Fold Change) and Y-axis to −log10(p-value). (C) Hierarchical clustering analysis sorted the control (pink) versus B_12_-deficient (green) group based on differential abundance of six metabolites (taurine, hypoxanthine, xanthine, glycocholic acid, neopterin, and chenodeoxycholic acid). Relative abundance scored from 4 (highest, red color) to −4 (lowest, blue). (D) MSEA plot with top 50 enriched metabolic pathways (vertical-axis) to which the 77 identified metabolites belong. The pathways are arranged in descending order of fold enrichment score (horizontal axis) where the highest is 6 (red color) and lowest is 0 (yellow color) (E) MetPA plot showing most enriched pathways with significance (-logP) values for each of the pathway as dots of red (high significance) or yellow (low significance). X-axis corresponds to pathway impact and Y-axis to -logP values. The size of the dot represents its impact value. (F) VIP score plot from PLS-DA analysis showing the top 20 differentially expressed metabolites in serum of control versus B_12_-deficient group scored from 1 to 2. Relative abundance is depicted with red (highest) and green (lowest) color. (G) Box plots showing normalized concentrations of individual metabolites following univariate analysis: taurine (p=0.002), xanthine (p=0.019) and hypoxanthine (p=0.000), chenodeoxycholic acid (p=0.063), neopterin (p=0.023), and glycocholic acid (p=0.027) in the sera of control (red) versus B_12_-deficient (green) groups.

Once we identified the most significant DEMs and major pathways to which these DEMs belonged to, we wanted to check the consistency of identified DEMs as most discriminant variables for classifying healthy controls versus B_12_-deficient subjects. For this purpose, we performed a PLS-DA analysis that helps in highlighting whether a metabolite is upregulated or downregulated in a group/sample by creating a latent structure, and the values of variable importance projection (VIP) score which represent the importance of the metabolite in the PLS-DA model (**Figure 2F**). The VIP score plot (threshold of >1.0) revealed that taurine had the maximum score with low abundance in B_12_-deficient samples versus controls (**Figure 2F**). The other metabolites that were identified in volcano plot i.e., xanthine, hypoxanthine, chenodeoxycholic acid, neopterin, and glycocholic acid also came up in PLS-DA plot, suggesting the consistency of these metabolites as important DEMs in controls versus B_12_-deficient subjects. Further, we performed univariate analysis (t-test) on individual DEMs to determine the significant difference in the abundance of each metabolite between the two groups. Based on the analysis, abundance of taurine (*p*=0.002), xanthine (*p*=0.019) and hypoxanthine (*p*=0.000) was significantly lower whereas the levels of chenodeoxycholic acid (*p*= 0.063), neopterin (*p*= 0.023), and glycocholic acid (*p*= 0.027) was significantly higher in sera of B_12_-deficient subjects (green bars) compared to healthy controls (pink bars) (**Figure 2G**).

Metabolites that belong to the same pathway tend to work in coherence. To this end, we subjected the metabolite data to Pearson’s correlation matrix analysis to reveal any correlation that might exist between the 77 identified metabolites or between 21 study subjects (**Figure S1A-B**). Between the two cohorts, metabolites such as taurine, xanthine, and hypoxanthine were positively correlated (red color) to each other and negatively correlated (blue color) to chenodeoxycholic acid, neopterin, and glycocholic acid (**Figure S1A**). Moreover, there was a high positive correlation observed between all the essential amino acids. This suggests a strong inter-relationship between these metabolites which could be expected as these belong to same metabolic pathway such as amino acid biosynthesis. Pearson’s correlation matrix analysis on the different cohort subjects, however, revealed no significant trends (**Figure S1B**), suggesting no inter-relationship or correlation between the samples, which negates the possibility of any biases in the sample workflow.

Taken together, these multiple lines of evidence suggest that taurine, hypoxanthine, xanthine, chenodeoxycholic acid, neopterin, and glycocholic acid are the most significant DEMs in the sera of healthy controls versus B_12_-deficient subjects. Pathway enrichment analysis further confirmed that the alteration in taurine and hypoxanthine metabolic pathway is strongly associated with B_12_ deficiency.

### Selection and identification of metabolite and/or metabolite ratio as biomarker

To identify the best metabolite and/or metabolites ratio that could serve as a sensitive biomarker for prediction of B_12_ deficiency, we subjected the data to two statistical analysis tools: Partial least squares discriminant-analysis (PLS-DA) (**Figure 3A and 3E**) and Random forest (RF) analysis (**Figure 3C and 3G**). Multiple statistical models generated by these analyses were validated and compared for their ability to identify the metabolite or metabolites ratio which can serve as the best biomarker to predict B_12_ deficiency. All models generated by PLS-DA or RF were validated using Receiver Operating Characteristic (ROC) analysis, in which Area Under the Curve (AUC) score was used to monitor the sensitivity and specificity of a model (variable) in predicting the B_12_ deficiency. Although both are predictive modelling tools, PLS-DA analysis has a tendency to overfit even on completely random data as compared to RF analysis. Thus, the quality of the models was further assessed using Monte-Carlo cross validation (MCCV) to create ROC curve for every model generated from both PLS-DA and RF analysis. These models use a combination of the most important features to build classification models, ranging from a minimum of 2 to a maximum of 100. Since MCCV uses defined sub-sampling, 2/3 of the samples were used to evaluate the feature importance and 1/3 of the samples were used for validation. This iterative procedure was used to calculate the performance (AUC) and confidence interval of each model and the one with AUC closest to 1 with low variability (CI) was considered to be the best model. The software gave output in the form of ROC curves of top 6 models, referred to as variables, based on the CV performance. we used the most significant DEMs (**Figure 3A & C**) or metabolite ratio (**Figure 3E & G**) as top features to generate best 6 models for prediction of B_12_ deficiency. Note that the nomenclature of models (referred to as variables, hereinafter) is representative of the number of features used to create the model. **Figure 3B, D, F**, and **H** represent the ROC curve for the top 6 models obtained following PLS-DA and RF analysis, whereas the model numbers 1, 2, 3, 4, 5 and 6 represent the variables (Var.) 3 (red), 5 (green), 10 (blue), 20 (cyan), 28 (pink) and 77 (yellow), respectively, signifying that model 1 was created using 2 metabolites of top importance, whereas model 6 used top 77 metabolites.

**FIGURE 3:**
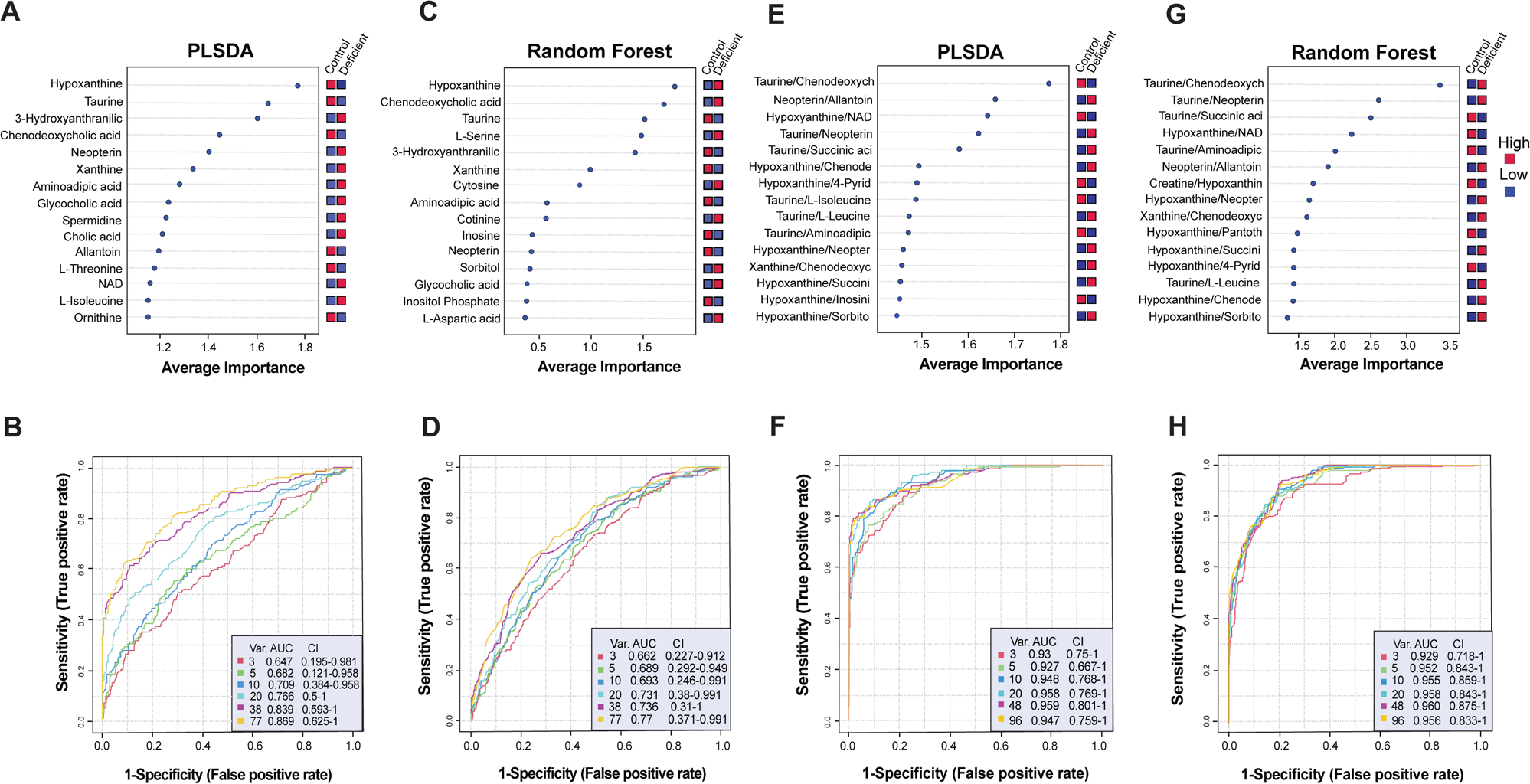
Selection and identification of metabolite and/or metabolite ratio as a biomarker. The top 6 predictive models (Var.) generated by various multivariant analyses were compared for their performance as metabolite biomarker predictors for B_12_ deficiency using ROC-AUC curves based on the MCCV method. ROC-AUC curve for (A) PLS-DA and (C) RF models using singular metabolites as features. ROC-AUC curve for € PLS-DA and (G) RF models using abundance ratio of metabolite pairs as features. Feature ranking plot for (B) PLS-DA and (D) RF models representing the top 15 metabolites arranged in descending value of average importance score. The average importance scores range from 1 to 2 for PLS-DA and 0 to 2 for RF. Feature ranking plot for (F) PLS-DA and (H) RF models representing top 15 abundance ratio of metabolite pairs arranged in descending value of average importance score. The average importance score ranges from 1 to 2 for PLS-DA and 1 to 4 for RF. In all the feature ranking plots the relative abundance of each feature between the control and B_12_-deficient group was graded with red and blue colors representing high and low abundance, respectively.

Both PLS-DA (**Figure 3A**) and RF (**Figure 3C**) analysis, using singular metabolites as features, showed that models with more than 20 metabolites (38 and 77) have high AUC (> 7) and tight CI, suggesting their potential to be better models, compared to those with fewer than 20 metabolites. A higher score suggests better predictive ability of a model to identify the B_12_-deficient state. The feature ranking plot for both PLS-DA (**Figure 3B**) and RF (**Figure 3D**) analysis showed the top 15 metabolites arranged in descending order of average importance scores contributing to the model accuracy. The average importance scores of hypoxanthine and taurine were among the top three metabolites in both analyses, with hypoxanthine having the maximum score. Both models showed lower (blue) abundance of taurine in B_12_-deficient cohort, but the same was not true for hypoxanthine. This was consistent with PLS-DA analysis done in **Figure 2F**. It is important to note that (a) 7 of 15 top metabolites were different between the models generated by PLS-DA and RF and (b) the individual average importance score for the 8 identical metabolites varied in the two analyses. This suggested that both analyses work on independent algorithms and there was no bias in the selection of hypoxanthine and taurine as top metabolite biomarkers for predicting B_12_ deficiency.

Next, we investigated whether abundance ratios of metabolite pairs could increase the sensitivity of PLS-DA and RF models to detect B_12_ deficiency (**Figure 3C**, **3D**). Ratios of all possible metabolite pairs were computed, and top ranked ratios (based on p values) and top 20 were included for biomarker analysis. Using abundance ratios of metabolite pair as a feature, both PLS-DA (**Figure 3E**) and RF (**Figure 3G**) models showed that all the top 6 models have high AUC (> 9) and high CI which were comparable, suggesting any model with more than 3 features was a good model with high specificity and sensitively but high variability (scattered CI) as well. One-to-one comparison of AUC and CI scores for both the PLS-DA and RF models based on the abundance ratios of metabolite pair versus singular metabolites revealed that the former can serve as better biomarkers in predicting B_12_ deficiency. The feature ranking plot for models in **Figure 3F** and **Figure 3H** listed 13 identical sets of metabolite pairs with taurine/chenodeoxycholic acid gaining the highest average importance score in both (**Figure 3G-H**). The abundance for taurine/chenodeoxycholic acid ratio however was reversed in the two models, being low (blue) in PLS-DA and high (red) in RF for B_12_-deficient group (**Figure 3E, 3G**). It is important to note that this analysis was consistent with the previous analysis shown in **Figure 2** (PCA, volcano plot, PLS-DA and univariate analysis).

Together, results suggest that out of the metabolites identified to be differentially expressed between healthy controls and B_12_-deficient group taurine, hypoxanthine and the ratio of taurine/chenodeoxycholic acid could serve as biomarkers for B_12_ deficiency.

### Comparison of the abilities of taurine, hypoxanthine and taurine/chenodeoxycholic acid ratio to predict B_12_-deficient state

We performed ROC analysis to further characterise the predictive ability of taurine alone, hypoxanthine and taurine/chenodeoxycholic acid ratio, which were shortlisted from previous PLS-DA and RF analysis. The sensitivity and significance of taurine, hypoxanthine and taurine/chenodeoxycholic acid in predicting B_12_ deficiency is represented using AUC score from ROC analysis (**Figures 4A-C**). The scaled concentration of the indicated metabolites are shown in **Figures 4D-F**. This analysis showed that AUC for taurine/chenodeoxycholic abundance ratio was 1, which is equivalent to being a perfect diagnostic biomarker (**Figure 4C**). Furthermore, the AUC and *p*-values for taurine/chenodeoxycholic acid ratio were the lowest (*p*-value=5.3193E-7) in comparison to hypoxanthine (AUC = 0.885, *p*-value =7.0513E-4) and taurine alone (AUC = 0.885, *p*-value =0.002), suggesting that taurine/chenodeoxycholic ratio was the best variable as a biomarker to predict B_12_ deficiency compared to others. Between taurine and hypoxanthine, the AUC scores were comparable, but hypoxanthine was significant in differentiating the two groups because of lower *p*-value.

**FIGURE 4:**
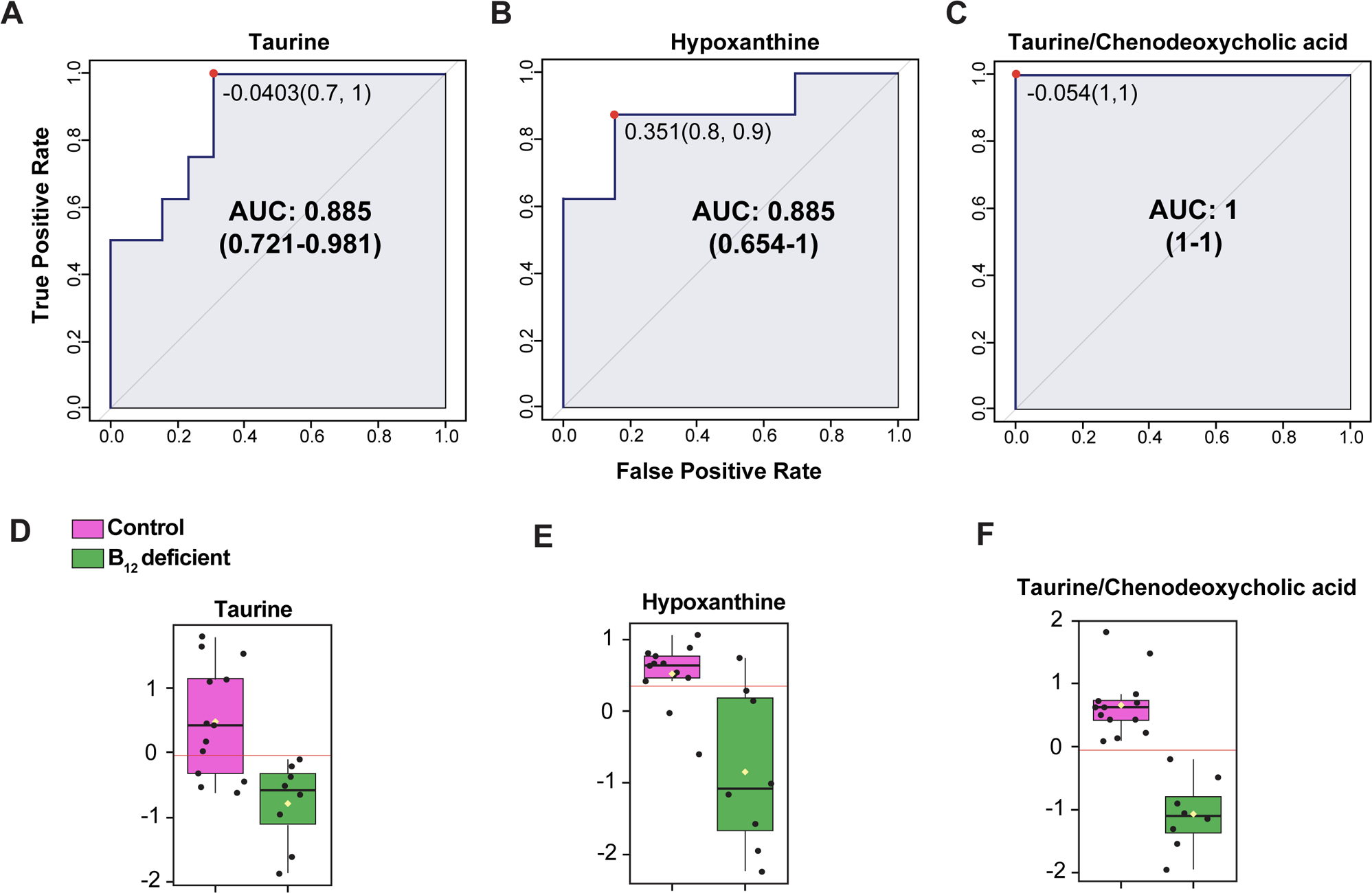
Comparison of the abilities of taurine, hypoxanthine and taurine/chenodeoxycholic acid ratio to predict B_12_-deficient state. ROC-AUC curve showing performance of (A) taurine, (B) hypoxanthine and (C) taurine/chenodeoxycholic acid ratio as biomarker to predict B_12_ deficiency based on AUC (sensitivity, specificity) and CI (variability) values. Each ROC curve is a plot between false positive rate (x-axis) and true positive rate (y-axis). Box plots showing normalized concentration of (D) taurin€(E) hypoxanthine and (F) taurine/chenodeoxycholic acid ratio between control (pink) versus B_12_-deficient (green) group. Each dot represents a sample. Y-axis represents fold change values. P value <0.05.

These results suggest that serum taurine/chenodeoxycholic acid abundance ratio can serve as a diagnostic biomarker for predicting B_12_ deficiency with high specificity and sensitivity.

To further test the ability of RF using taurine alone or and in combination with other metabolites as biomarker to predict B_12_ deficiency, we trained a RF model on train data using cross validation and predicted on the test data. For unbiased assessment, equal number of samples (n=4/group) were randomly selected from control and B_12_-deficient group as hold-out samples. These samples were not used for fitting process in the model but used as testing samples. The rest of the samples were used as training samples to predict B_12_ deficiency. We compared predictive ability of taurine alone, taurine and hypoxanthine, and ratio of taurine/chenodeoxycholic acid using AUC score (ROC analysis), predicted class probabilities, and cross validation (CV) prediction (**Figure 5**). Amongst these model (**Figure 5A, 5C, 5E**) comparisons, taurine/chenodeoxycholic acid showed the highest margin of separation between the control (empty grey circles, left edge of x-axis) and B_12_-deficient (filled grey circles, right edge of x-axis) group in training set, (**Figure 5E**). Also, the hold-out samples from both groups (control = empty red circles, B_12_-deficient = red filled circles) fit perfectly well with the corresponding group in testing data set. Moreover the ROC-AUC curve showed that taurine/chenodeoxycholic abundance ratio had the highest accuracy (AUC CV=1, AUC holdout =1, **Figure 5F**) in predicting B_12_ deficiency compared to taurine alone (AUC CV = 0.665, AUC holdout=0.938, **Figure 5B**) or hypoxanthine (AUC CV= 0.809, holdout=0.938, **Figure 5D).** Overall, this analysis was consistent with previous RF analysis, suggesting towards great potential of taurine/chenodeoxycholic acid to serve as serum biomarker for predicting B_12_ deficiency.

**FIGURE 5:**
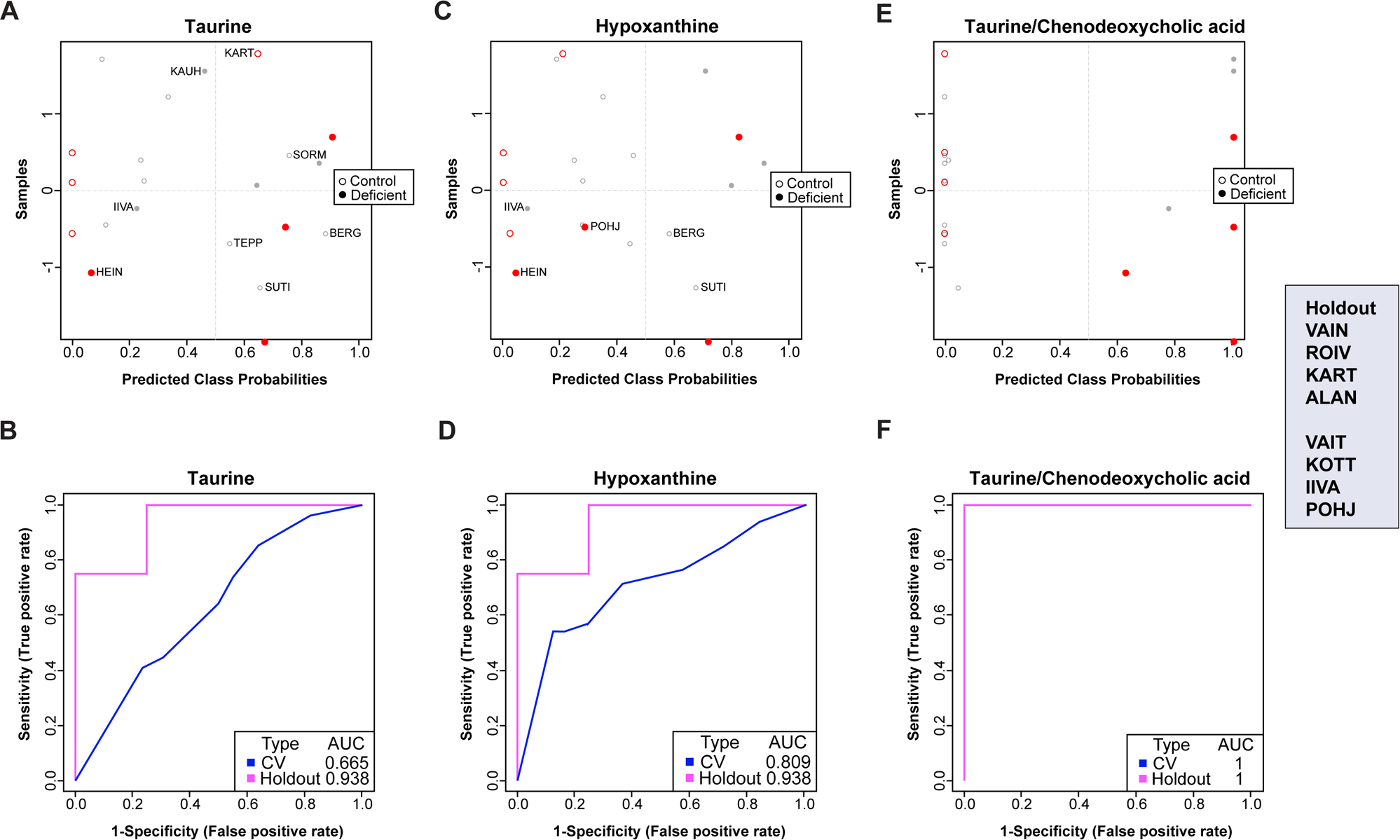
Statistical Model to test predictive ability of taurine alone and in combination as biomarker. Random forest was used as a model to test the predictive abilities of taurine, taurine and hypoxanthine together, and taurine/chenodeoxycholic acid ratio to predict B_12_ deficiency. Predicted class probability plot for (A) taurine, (B) taurine and hypoxanthine together, and (C) taurine/ chenodeoxycholic acid ratio showing the classification accuracy of each factor to differentiate between control (grey dots) and B_12_-deficient (red dots) samples. The solid dots are training data sets and the empty dots are test data sets. ROC-AUC curve analysis showing cross-validation (pink) and hold-out (blue) scores to determine the performance of (D) tau€e, (E) taurine and hypoxanthine, and (F) taurine/chenodeoxycholic acid ratio as a biomarker to predict B_12_ deficiency. Each ROC curve is a plot between the false positive rate (specificity) on the x-axis and true positive rate (sensitivity) on the y-axis.

### Metabo-transcriptomic network analysis linked B_12_-dependent reactions with taurine/chenodeoxycholic acid

We performed a network analysis of differentially expressed genes and metabolites between controls and B_12_-deficient livers in a mouse model of B_12_ deficiency reported previously by us.(28) Liver is a suitable tissue to investigate effects of B_12_ deficiency since it is one of the principal site of B_12_ storage, and we demonstrated earlier that B_12_ deficiency compromises its functions.(28) In the cells, B_12_ is known thus far to be converted into two cofactors (methyl-B_12_ and adenosyl-B_12_), which are required for the functioning of two known enzymes, methionine synthase and methyl-malonyl CoA mutase.(29, 30) Thus, we focused our attention on metabolic pathways that are interconnected with the B_12_-derived cofactor-dependent reactions such as Krebs cycle, amino acid metabolism, urea cycle, and nucleotide metabolism.

The network visualization of differentially expressed transcriptome showed that transcripts encoding the enzymes that catalyze metabolite conversions in these pathways were overall downregulated (in blue), except for the Krebs cycle, in which expression of 5 out of 9 enzymes was upregulated (in red) (**Figure 6**). This upregulation in the expression levels of Krebs cycle enzymes could be linked to decreased activity of methyl-malonyl CoA mutase (Mut), which is dependent on the adenosyl-B_12_ for its activity. Mut catalyzes the synthesis of Succinyl-CoA, an intermediate in the Krebs cycle that plays a critical role in providing protons for the OXPHOS system, and thus, energy production in the cells. B_12_ deficiency leads to an energy deficit in the cells, and consequently likely, a compensatory increase in the expression levels of enzymes in the Krebs cycle. However, no reactions surrounding the adenosyl-B_12_-dependent Mut enzyme and Krebs cycle could relate to known taurine biosynthetic machinery in B_12_-deficient cells.

**FIGURE 6:**
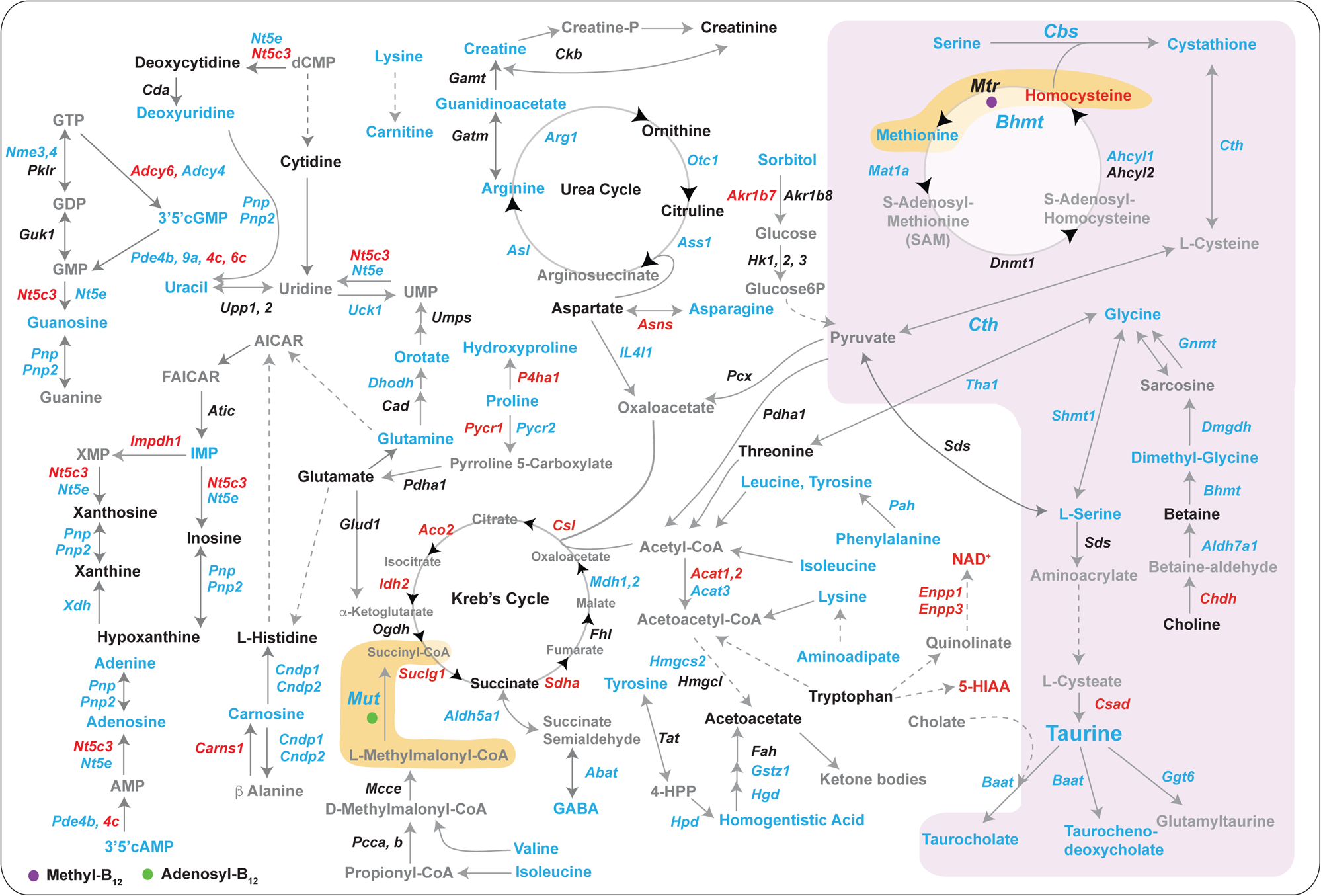
Metabo-transcriptomic network analysis links B_12_ dependent reactions with taurine/chenodeoxycholic acid. Network analysis showing the differentially expressed genes and metabolites between controls and B_12_-deficient livers in a mouse model of B_12_ deficiency reported previously(25). The network shows interactions between enzymes (italics font) and metabolites (normal font) across various metabolic pathways in the liver such as Krebs cycle, urea cycle, amino acid metabolism, nucleotide metabolism, etc. The arrows represent the direction of the reaction. The downregulation and upregulation of enzyme transcript or metabolite concentrations are represented by blue and red color, respectively.

An analysis of reactions surrounding methionine synthase (Mtr), the second enzyme that is dependent on the methyl-B_12_ as a cofactor, showed that the concentrations of methionine, the downstream product, were decreased while concentrations of its precursor, homocysteine, were increased (**Figure 6**). Expression levels of the enzymes in the methionine cycle were either not affected or were decreased. The methionine cycle is linked to cysteine synthesis in the cells and through a relay of changes, to taurine biosynthesis. Most of the enzymes and their downstream products in this pathway were downregulated, consequently leading to deficiency of multiple metabolites in taurine metabolic pathway (taurine, taurocholate, tauro-chenodeoxycholate) (**Figure 6**). The expression levels of the enzyme, *Csad*, that catalyzes the rate limiting step in taurine biosynthesis, was increased likely as a compensatory mechanism due to deficiency of taurine (**Figure 6**).

Further analysis of gene-metabolite networks interconnected with B_12_-dependent reactions showed that gene expression of enzymes and metabolite intermediates in the urea cycle were downregulated. In the amino acid metabolism pathway, barring tryptophan metabolite, HIAA and NAD^+^ pathways, all enzyme expressions and metabolite intermediates were downregulated. In the nucleotide metabolism pathways, metabolite intermediates were either downregulated or not affected, and apart from a few enzymes, most of the enzyme expressions were downregulated.

Together, these integrated metabolomic and transcriptomic analyses in the WT and B_12_-deficient liver samples revealed global downregulation of metabolic networks upon B_12_ deficiency and identified a hitherto unanticipated connectivity between B_12_-dependent reactions and taurine metabolism.

## DISCUSSION

By using metabolomic analysis of serum from controls and B_12_-deficient subjects, we were able to identify that a ratio of taurine/chenodeoxycholic acid levels can serve as a biomarker of, difficult to detect, B_12_ deficiency. The quantitative metabolomic analysis of 77 relevant metabolites in the sera of B_12_-deficient patients revealed that most of the metabolites were downregulated and are involved in metabolism of amino acids, betaine, glutathione, bile acid, and purines (**Figure 2**). Metabolite set enrichment analysis on the perturbed metabolite profiles showed alterations in the metabolic pathways associated with amino acid and methionine metabolism (**Figure 1**). Downregulation in methionine levels in this metabolome is consistent with the role of B_12_ as an essential cofactor of methionine synthase, while homocysteine accumulated from the dysfunction of methionine synthase was 1.8-fold elevated. Furthermore, univariate analysis of the B_12_-deficient metabolome identified a differential abundance of taurine, hypoxanthine, and xanthine between the two groups. The multivariate random forest (RF) analysis aimed towards identifying which metabolite(s) contributed to the separation of the two groups with higher specificity and sensitivity showed taurine/chenodeoxycholic ratio as the metabolic parameter that could separate the two groups with 99% accuracy. Thus, we propose taurine/chenodeoxycholic acid ratio as a potential biomarker of a B_12_-deficient state in humans.

Previous studies have characterized the human serum metabolome in B_12_-deficient subjects in an attempt to reveal connections between B_12_-deficient state and serum metabolic markers. Alex et al., performed metabolomic profiles in sera of Chilean older adults with subclinical borderline B_12_ deficiency (defined by serum B_12_ <148 pmol/L, holotranscobalamin <35 pmol/L, tHcy >15 μmol/L, or MMA >271 nmol/L).(31) Although, this study showed perturbations in multiple metabolite such as acylcarnitine and plasmalogens Authors did not subject their data to downstream algorithms to identify potential biomarkers of B_12_ levels. Moreover, the previous study did not include a control group, whereas our study has a well-defined control group. Although, these studies provide evidence that serum metabolome is altered by B_12_ deficiency it was unknown whether any of the metabolites of set of metabolites could serve as a biomarker of B_12_-deficient state. Our study fills this gap in our knowledge and elucidates the effect of B_12_ deficiency on the cellular, metabolic and transcriptomic landscape of the cell using liver biopsies from a B_12_-deficient mouse model. Together, these studies pave a way towards better understanding of the cellular defects caused by B_12_ deficiency.

We acknowledge that our study has certain limitations. Firstly, the small sample size limits the statistical power of the RF models. Repeating the same study in a larger sample size may allow a greater number of metabolites to pass quality control for downstream analysis. Secondly, the current study population was only tested for B_12_ deficiency, which does not rule out the possibility of deficiency of other vitamins or nutrients in the study population. These and other questions will need to be addressed in future studies.

Vitamin B_12_ deficiency leads to perturbed levels of taurine, hypoxanthine, xanthine, chenodeoxycholic acid, neopterin, and glycocholic acid. We show that taurine levels alone and taurine/chenodeoxycholic acid ratio are promising candidates for serum metabolite-based biomarkers to identify B_12_ deficiency. The two critical metabolites identified in this study regulated by B_12_, taurine and chenodeoxycholic acid, belong to the taurine metabolic pathway. Taurine metabolism gets compromised with age and leads to taurine deficiency in humans, however, the cause of this deficiency is unknown(24). The present study identifies vitamin B_12_ as the very first upstream regulator of taurine metabolism in aged humans and illustrates the transcriptomic and metabolomic changes through which B_12_ regulates this process. These results are significant given that taurine deficiency has recently been shown to be a driver of aging in diverse species, and is associated with poor health in humans. This study paves a way for future clinical work to streamline diagnostic tools to detect B_12_ deficiency through a simple blood test and perhaps other age-associated diseases.

## Acknowledgements

We thank Research Support Facility staff at Sanger Institute and National Institute of Immunology especially Dr. P. Nagarajan for assistance with animal experiments.

## Authorship

All authors have seen and approved the manuscript

## MATERIAL AND METHODS

### Chemicals and reagents

All the metabolite standards, ammonium formate, ammonium acetate and ammonium hydroxide were obtained from Sigma-Aldrich (Helsinki, Finland). Formic acid (FA), 2-proponol, acetonitrile (ACN), and methanol (all HiPerSolv CHROMANORM, HPLC grade, BDH Prolabo) were purchased from VWR International (Helsinki, Finland). Isotopically labelled internal standards were obtained from Cambridge Isotope Laboratory. Inc., USA (Ordered from Euriso-Top, France). Deionized Milli-Q water up to a resistivity of 18 MΩ L cm was purified with a purification system (Barnstead EASYpure RoDi ultrapure water purification system, Thermo scientific, Ohio, USA).

### Metabolite extraction protocol

The working calibration solutions were prepared in 96-well plate by serial dilution of the stock calibration mix using Hamilton’s MICROLAB® STAR line (Hamilton, Bonaduz AG, Switzerland) liquid handling robot system. Starting from a stock solution mix, 10 additional lower working solutions were prepared using water as the diluent to build the calibration curves.

### Clinical serum samples

Clinical samples used for assessing the changes in vitamin B_12_ levels and metabolites in blood were obtained from the Kuopio Ischaemic Heart Disease Risk Factor Study (KIHD study), a population-based cohort study described previously (25, 32), and were donated by J. Kauhanen and T. Nurmi (University of Eastern Finland, Kuopio, Finland). Ten microliters of labelled internal standard mixture was added to 100 μL of serum sample. Metabolites were extracted by adding 4 parts (1:4, sample: extraction solvent) of the 100% ACN + 1% FA solvent. The collected extracts were dispensed in OstroTM 96-well plate (Waters Corporation, Milford, USA) and filtered by applying vacuum at a delta pressure of 300-400 mbar for 2.5 min on robot’s vacuum station. This resulted a cleaner extract to the 96-well collection plate, which was placed under the OstroTM plate. The collection plate was sealed with the cap map and placed in auto-sampler of the LC system for the injection.

### Instrumentation and analytical conditions

Sample analysis was performed on an ACQUITY UPLC-MS/MS system (Waters Corporation, Milford, MA, USA). The auto-sampler was set at 5°C, and the column, 2.1 × 100 mm Acquity 1.7um BEH amide HILIC column (Waters Corporation, Milford, MA, USA), temperature was maintained at 45°C. The total run time is 14.5 min including 2.5 min of equilibration step at a flow rate of 600 μL/min. Initially the gradient started with a 2.5 min isocratic step at 100% mobile phase B (ACN/ H2O, 90/10 (v/v), 20 mM ammonium formate, pH at 3), and then rising to 100% mobile phase A (ACN/H2O, 50/50 (v/v), ammonium formate, pH at 3) over the next 10 min and maintained for 2min at 100% A and finally equilibrated to the initial conditions for 2.5 min. An injection volume of 5 μL of sample extract was used and two cycles of 300 μL of strong wash (methanol/isopropanol/ACN/H2O, 25/25/25/25, 0.5% FA) and 900 μL of weak wash (methanol/isopropanol/ACN/H2O, 25/25/25/25, 0.5% ammonium hydroxide) and in addition 2 min of seal wash (90/10, methanol/H2O) were carried out. The auto-sampler was used to perform partial loop with needle overfill injections for the samples and standards.

The detection system, a Xevo® TQ-S tandem triple quadrupole mass spectrometer (Waters, Milford, MA, USA), was operated in both positive and negative polarities with a polarity switching time of 20 msec. Electro spray ionization (ESI) was chosen as the ionization mode with a capillary voltage at 0.6 KV in both polarities. The source temperature and desolvation temperature of 120°C and 650°C, respectively, were maintained constantly throughout the experiment. Declustering potential (DP) and collision energy (CE) were optimized for each compound. High pure nitrogen and argon gas were used as desolvation gas (1000 L/hr) and collision gas (0.15 ml/min), respectively. Multiple Reaction Monitoring (MRM) acquisition mode was selected for quantification of metabolites with individual span time of 0.1 sec given in their individual MRM channels. The dwell time was calculated automatically by the software based on the region of the retention time window, number of MRM functions and depending on the number of data points required to form the peak. MassLynx 4.1 software was used for data acquisition, data handling and instrument control. Data processing was done using TargetLynx software and metabolites were quantified by using labelled internal standards and external calibration curves.

### Data analysis using MetaboAnalyst 5.0 software and downstream analysis

The raw data was analyzed using MetaboAnalyst 5.0 software (https://www.metaboanalyst.ca/). (33, 34) Metabolite raw values were generalized log (glog) transformed and auto-scaled (mean-centered and divided by the standard deviation of each variable).(35) Missing values for any metabolites in the sample below the limit of detection were inputted with 1/5 of the minimum positive value for each variable. Unsupervised Principal component analysis (PCA) was done to differentially cluster the two groups.(36, 37) Hierarchical clustering and Pearson’s correlation analysis were also performed to cluster the metabolite and sample data in the form of a heatmap to easily identify patterns in metabolite concentrations across samples. Metabolite Set Enrichment Analyses (MSEA)(38) were performed on all metabolites with a VIP ≥1.5 that matched the database using the “Pathway-associated metabolite sets (SMPDB)” database in the MetaboAnalyst software. Pathway analysis was performed using the “Homo sapiens (KEGG(39, 40))” database in the MetaboAnalyst software. Interactive scatter plot with ‘Enrichment Factor’ as *x* axis and ‘−log_10_(*P*)’ as *y* axis was generated for functional analysis to show the significance of top 50 metabolic pathways involving the metabolites identified. The variable importance to projection (VIP) score for each metabolite was calculated to quantitatively represent metabolite feature importance in the model. A volcano plot scatterplot that shows statistical significance (-log10(p-value) versus magnitude of change (log 2-fold change) of metabolites. Metabolites that show significant (p ≤ 0.05) change (log 2-fold change ±0.5) are highlighted. Multivariate supervised Partial least squares discriminant analysis (PLS-DA) and Random-forest (RF) analysis were performed to assess the difference between the abundance of top metabolites or metabolite ratio between the two groups. The area under the curve (AUC) of the receiver operating characteristic (ROC) curve was also calculated for each metabolite to determine its predictive ability as a biomarker. The ROC curve is a plot of false positive rate (FPR) vs the true positive rate (TPR). The higher the AUC value, the better the measurements are at classifying between the two groups.

---

**FIGURE S1: Correlation analysis between metabolites and samples.** Pearson’s correlation matrix to identify highly correlated (A) metabolites and (B) samples in two groups. Correlation score ranged from 1 (highest, red) to −1 (lowest, blue).

