## Supplementary figures and images for "Taurine/chenodeoxycholic acid ratio as a circulating biomarker of insidious vitamin B_12_ deficiency in humans"

### Supplemental Figure 1

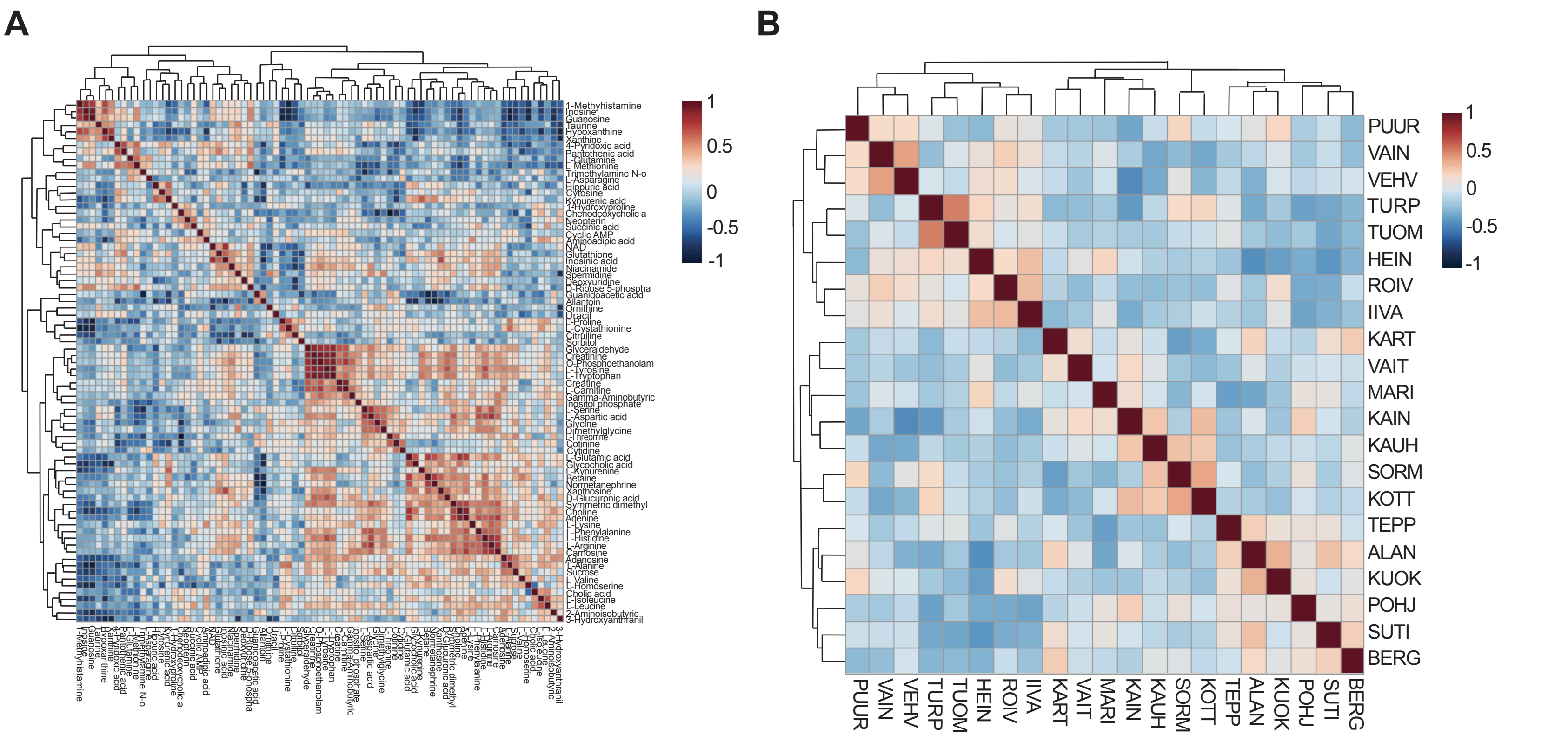
